# Anesthesia Lowers Spatial Frequency Preference in the Primary Visual Cortex

**DOI:** 10.1101/2025.09.29.679183

**Authors:** Jiahao Wu, Taisuke Yoneda, Kallum Robinson, Naotsugu Tsuchiya, Yumiko Yoshimura

## Abstract

This study examined the impact of general anesthesia on visual processing in the mouse primary visual cortex (V1). By directly comparing visual responses in the same layer 2/3 neurons between the awake and anesthetized states, we found that isoflurane anesthesia significantly shifts the preferred spatial frequency (SF) of neurons to lower values while leaving orientation tuning intact. This downward SF shift impaired the ability to encode fine visual details (high-SF) but not coarse information (low-SF). The shift was observed across all recorded cell types—including excitatory, somatostatin-expressing (SOM), and parvalbumin-expressing (PV) inhibitory neurons—though their respective magnitudes and underlying patterns were distinct. Furthermore, modulation of response gain and tuning sharpness was observed exclusively in SOM neurons. These results highlight the specific effects of anesthesia, which vary with visual feature and neuron type. These findings suggest that reduced sensory perception during anesthesia is linked to a degradation in spatial resolution at the cellular level.

## Introduction

General anesthesia is indispensable for the safe and humane execution of surgical, diagnostic, and therapeutic procedures. It produces an acute state of unconsciousness and unresponsiveness to external stimuli, representing one of the most extreme shifts in behavioral states. Extensive research has explored anesthesia’s effects and mechanisms, from identifying target molecules and neural pathways to understanding neuronal population dynamics and its relation with fluctuations in the level of consciousness ^1–5^. Previous studies in rodents have demonstrated that multiple types of general anesthesia induce spontaneous synchronous activity across local and inter-regional neuronal populations, as revealed by electrophysiological and optical imaging techniques ^6–10^.

A critical step in understanding the considerable reduction of sensory perception under anesthesia involves precisely characterizing whether anesthesia causes a uniform suppression of all sensory responses or selectively impairs specific features. It has been reported in humans and mice that while higher sensory cortical areas exhibit a significant reduction in their sensory responses under anesthesia, primary sensory cortices are less affected, even at similar anesthetic levels ^10,11^. The rodent primary visual cortex (V1), with its well-characterized receptive field properties, is thus a suitable area for investigating cell-level changes in primary cortical sensory responses induced by anesthesia. Previous studies have reported that orientation selectivity in V1 layer 2/3 is not affected under either urethane or isoflurane anesthesia ^12–14^. In contrast, size tuning and spatial selectivity in V1 neurons are reported to decrease under isoflurane anesthesia ^15,16^. These results suggest that the effects of anesthesia are visual parameter-specific. Spatial frequency (SF) tuning is a fundamental parameter for the extraction of object shape. Yet, how anesthesia modulates SF tuning has received little attention. Since SF preference is known to vary depending on a neuron’s receptive field position and its laminar location ^17–19^, isolating the effects of anesthesia requires tracking the same neurons across both awake and anesthetized conditions. However, this type of analysis has not been previously reported.

Previous studies in rodent V1 indicate that under isoflurane anesthesia, the response magnitude of excitatory neurons remains largely unchanged, while that of inhibitory neurons decreases ^16,20^. Additionally, visual responses are prolonged during anesthesia, possibly due to decreased inhibition ^16,21^. These findings suggest that inhibitory neurons are more sensitive to anesthesia than excitatory neurons. Cortical inhibitory neurons comprise multiple subtypes, each expressing different channel molecules sensitive to anesthesia ^5,22^. Furthermore, each subtype possesses distinct connectivity and functions ^23–26^. Consequently, the effect of anesthesia on sensory responses may differ across individual interneuron subtypes ^15^. However, these subtype-specific effects of anesthesia have not yet been systematically studied.

In this study, we directly compared visual responses recorded from the same V1 neurons in mice across anesthetized and awake states using two-photon Ca^2+^ imaging. To evaluate neuron type-specific effects of anesthesia, we utilized transgenic mice to label pan-inhibitory neurons, somatostatin-expressing (SOM) and parvalbumin-expressing (PV) interneurons. We found that anesthesia induced a shift in preferred SF to lower frequencies, irrespective of neuron subtype, and that the magnitude and pattern of SF shift were distinct for each subtype.

## Results

### Anesthesia Decreased Preferred Spatial Frequencies

To investigate how anesthesia affects visual response properties, we compared visual responses recorded from the same neurons in layer 2/3 of mouse V1 under both isoflurane-anesthetized (0.7–0.9%) and awake conditions. To this end, we used two-photon Ca^2+^ imaging. jGCaMP7b was expressed pan-neuronally using an AAV vector under the control of the synapsin promoter. Glutamatergic excitatory neurons were distinguished from GABAergic interneurons (pan-inhibitory neurons), which were labeled with red fluorescence (tdTomato) in Vgat-Cre;Ai14 mice (N = 9 mice; Fig. 1A). We recorded visual responses from tdTomato-negative, excitatory neurons, and tdTomato-positive, inhibitory neurons in response to full-field drifting sinusoidal grating stimuli, varying across five SFs and eight directions (Fig. 1B). For quantification, we selected neurons responsive to visual stimuli in both anesthetized and awake states (610 excitatory cells, 111 inhibitory cells, Table S1).

**Figure 1.**
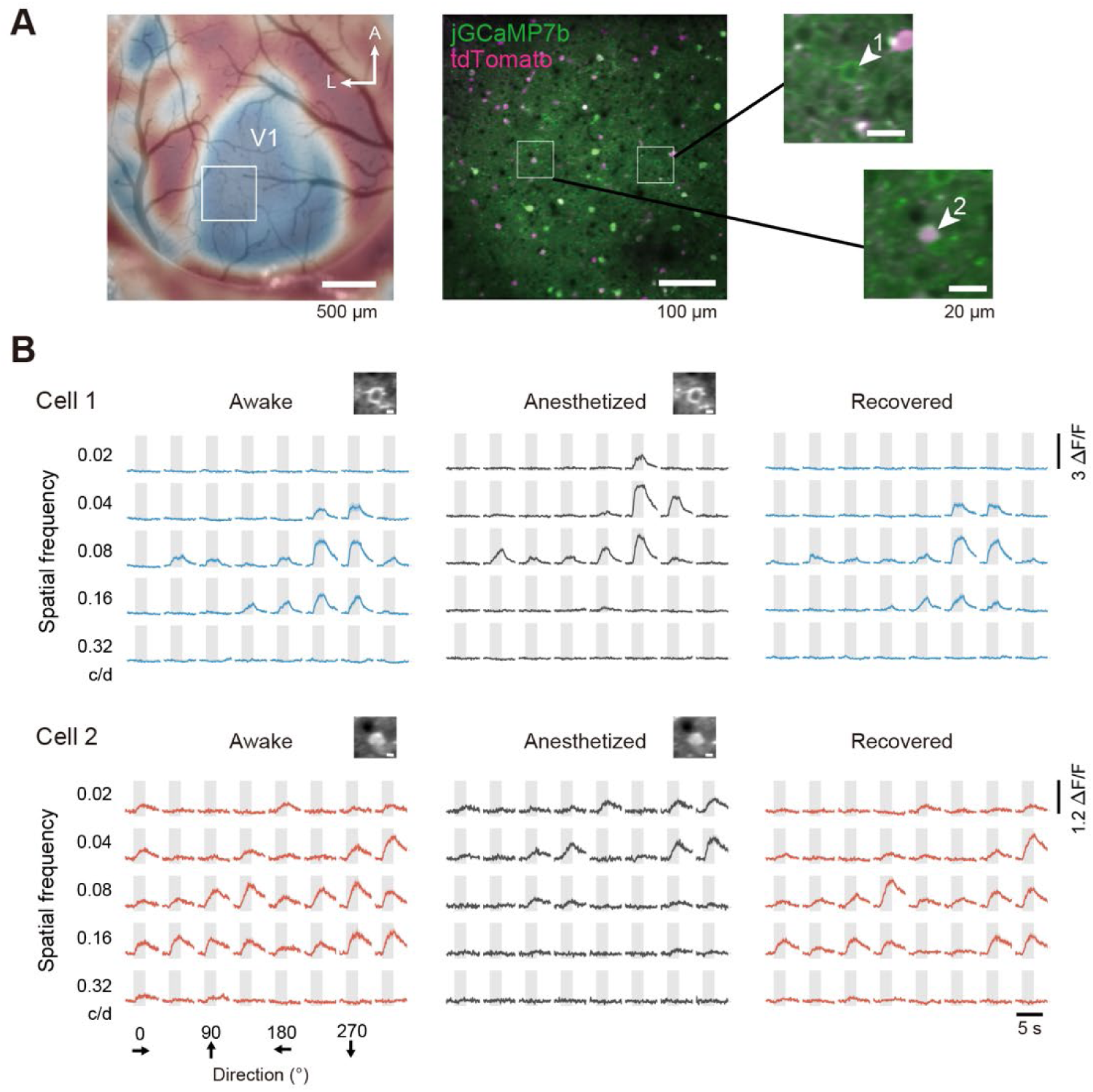
Two-photon Ca^2+^ imaging of the same neurons in the anesthetized and awake states. (A) The primary visual cortex (V1) was first mapped by wide-field Ca²⁺ imaging (left), followed by two-photon cellular-level imaging (middle). On the right, higher-magnification images show jGCaMP7b-expressing neurons without (cell 1) and with tdTomato labeling (cell 2) (B) Representative visual responses of a tdTomato-negative excitatory (cell 1) and a tdTomato-positive inhibitory neuron (cell 2) in the awake (left), anesthetized (middle) and recovery (2nd awake, right) conditions (mean ± SEM). Responses (ΔF/F) were evoked by visual stimuli with various spatial frequencies (1-octave steps, shown on the left) and directions (45°-steps, shown at the bottom). Each gray box indicates the stimulus presentation period (2 s). Top, mean image of the recorded neurons in each condition.

In the excitatory neurons, anesthesia decreased preferred SFs by 0.15 octaves (linear mixed-effects model, 95% CI [0.09, 0.20], p = 6×10^−8^; Fig. 2A). The differences in preferred SFs were statistically significant (median: anesthetized 0.059 c/d, awake 0.074 c/d; p = 3×10^−25^, Kolmogorov–Smirnov test, n = 610 cells; Fig. 2A). However, preferred orientation did not significantly change between anesthesia and wakefulness (p > 0.1, Kuiper two-sample test, Fig. S1), indicating that anesthesia specifically modified SF preference. To rule out order effects, visual responses were also recorded in five of nine mice after full recovery from anesthesia (>4h in the home cage; Figs. 1B and 2B). Preferred SFs did not differ between the initial awake session and the recovered condition (second awake session) (p = 0.97, Kolmogorov–Smirnov test, n = 376; Fig. 2B). Thus, the anesthesia-induced changes in SF tuning were not due to the recording order or representational drift ^27,28^.

**Figure 2.**
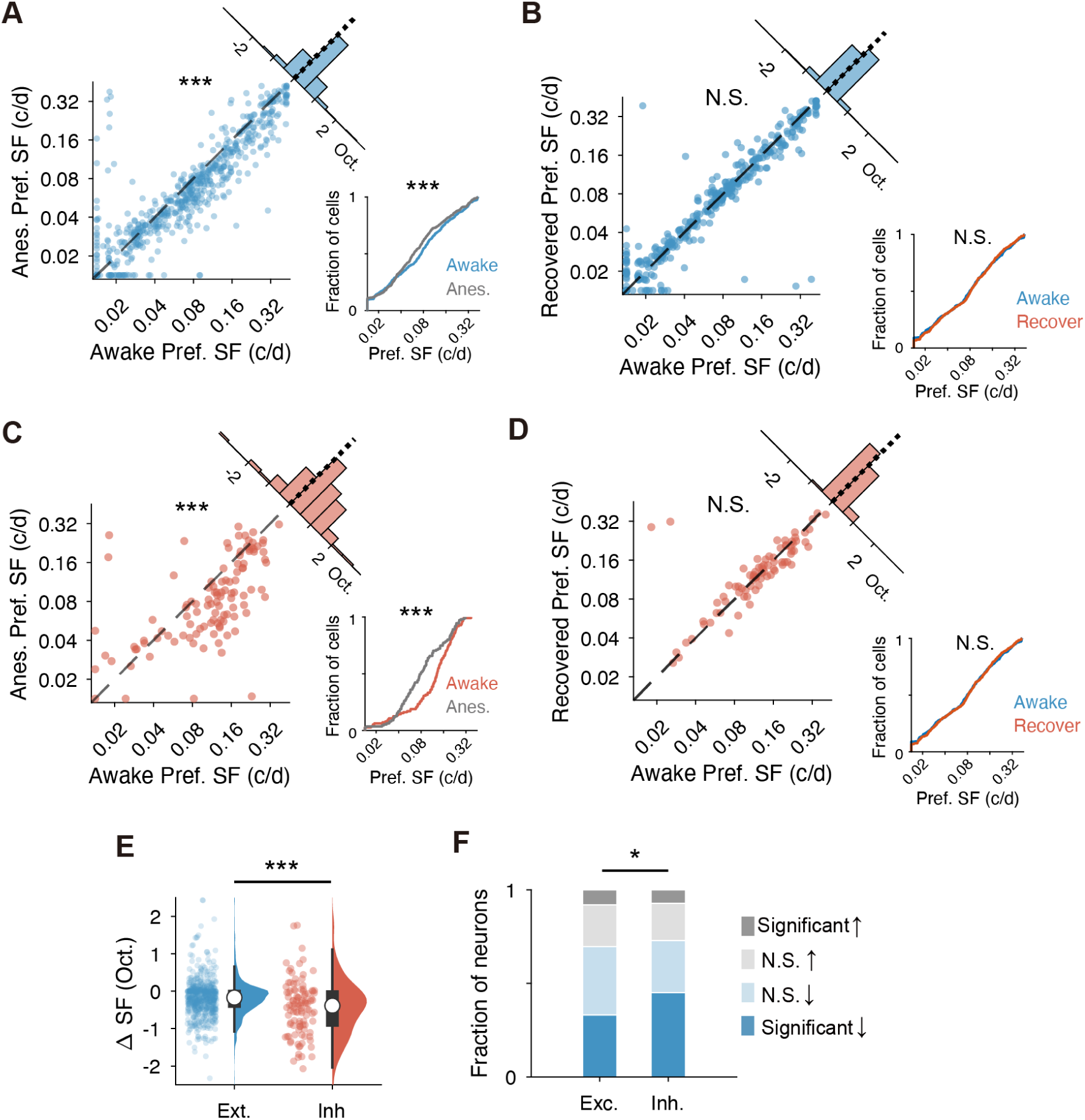
Anesthetize-induced downward shift of the preferred spatial frequency. (A) Preferred SF of individual neurons in the anesthetized condition (y-axis) is plotted against that in the awake condition (x-axis) for tdTomato-negative excitatory neurons (n = 610 cells, N = 9 mice). Linear mixed model; ***p < 0.001; N.S., not significant. The corner histogram shows the distribution of the pairwise difference in preferred SF between the two conditions. Bottom right: cumulative distributions of preferred SFs in each condition. Kolmogorov–Smirnov test; ***p < 0.001. (B) Preferred SF of excitatory neurons in the recovered condition from the anesthesia (2^nd^ awake, y-axis) is plotted against that in the 1^st^ awake condition (x-axis) (n = 376 cells, N = 5 mice). (C–D) Same plots for tdTomato-positive, inhibitory neurons (C, n = 111 cells, N = 9 mice; D, n = 80 cells, N = 5 mice). (E) Box and raincloud plots comparing the anesthesia-induced changes in preferred SFs for individual neurons between excitatory (Ext.) and inhibitory neurons (Inh.). ΔSF indicates the difference in preferred SF (from the awake state to the anesthetized state) in each neuron. Wilcoxon signed-rank test; ***p < 0.001. (F) Fractions of excitatory and inhibitory neurons exhibiting a significantly decreased (blue), non-significantly (N.S.) changed (light blue or light gray), or significantly increased (gray) preferred SF under anesthesia. Comparison of fractions between excitatory and inhibitory neurons. Chi-square test; *p < 0.05.

Next, we examined whether similar anesthetic effects occurred in inhibitory neurons. On average, preferred SFs in inhibitory neurons decreased by 0.42 octaves under anesthesia compared to wakefulness (linear mixed-effects model, 95% CI [0.24, 0.60], p = 9×10^−6^; Fig. 2C). This decrease was statistically significant (median: anesthetized 0.079 c/d, awake 0.13 c/d; p = 1×10^−5^, Kolmogorov–Smirnov test, n = 111; Fig. 2C). Similar to excitatory neurons, the preferred SFs did not differ between the initial awake session and the second awake session (p = 0.81, Kolmogorov–Smirnov test, n = 80; Fig. 2D). Inhibitory neurons showed a significantly larger anesthesia-induced shift in preferred SFs than excitatory neurons (p = 7×10^−5^, Wilcoxon signed-rank test; Fig. 2E). A significant anesthesia-induced downward shift in preferred SFs was observed in 34% of excitatory neurons and 45% of inhibitory neurons (bootstrap analysis, Fig. 2F, Fig. S2). The proportion for inhibitory neurons was significantly greater than that for excitatory neurons (p = 0.01, Chi-square test, Fig. 2F). These results suggest that while anesthesia modulates the SF preference of both excitatory and inhibitory neurons, the resulting downward shift of preferred SFs is more pronounced in inhibitory than in excitatory neurons.

### Anesthetic Effects on the Neuronal Population Coding

To investigate how anesthesia affects the population coding of visual stimuli, we performed a neural decoding analysis using a Bayesian decoder. The decoder was trained on visual responses from excitatory neuron populations in the awake state and subsequently applied to decode visual stimuli from neuronal activity under both awake and anesthetized conditions.

We first evaluated the information carried by population responses by measuring the orientation decoding error. While orientation decoding errors for low-SF stimuli (≤ 0.04 c/d) were comparable between the two states, errors for high-SF stimuli (> 0.04 c/d) were significantly increased under anesthesia (Fig. 3A). This finding suggests that anesthesia selectively degrades the neural information related to high-SF stimuli.

**Figure 3.**
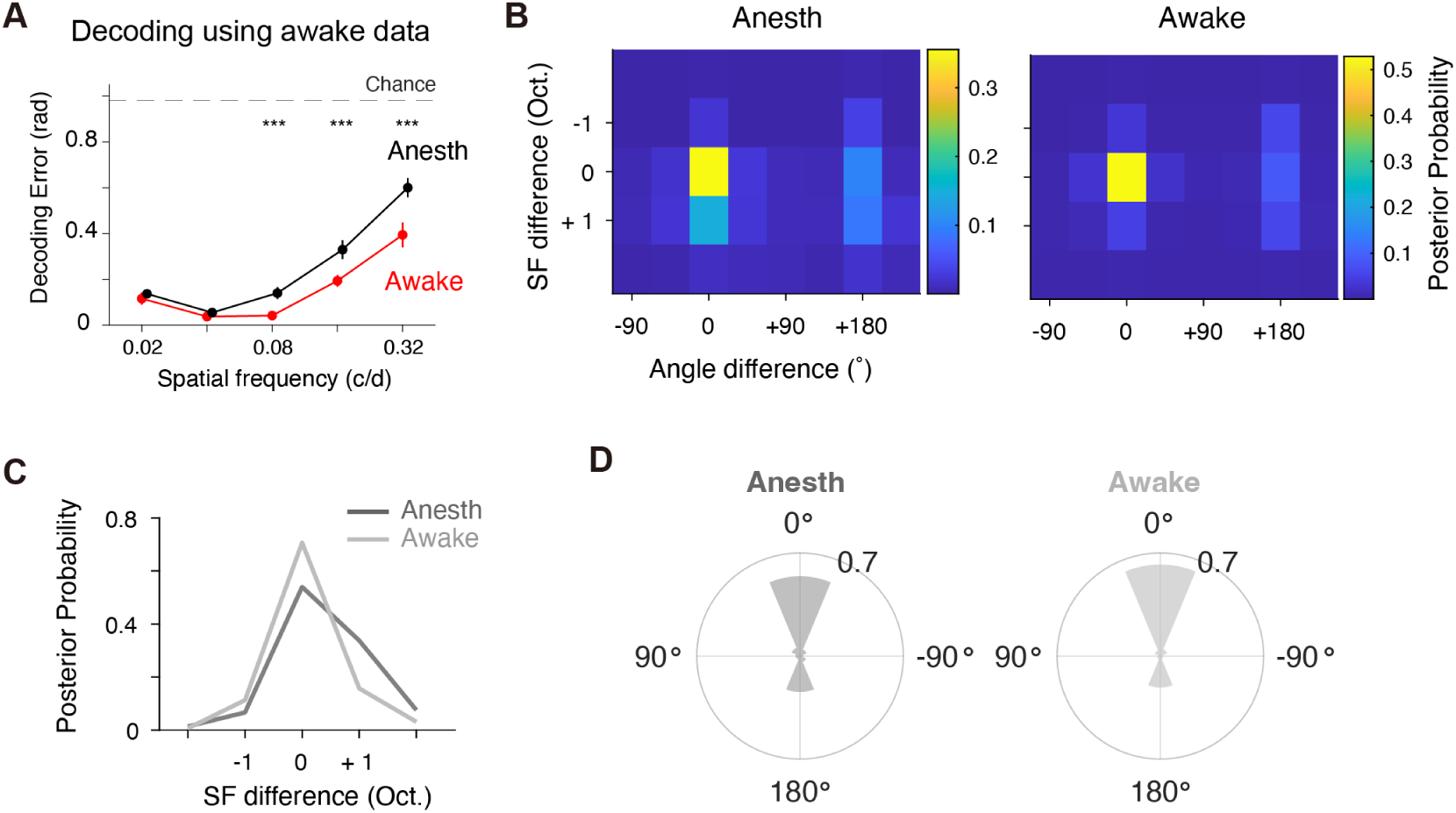
Effects of anesthesia on neuronal population coding. (A–D) A decoder was trained using visual responses in excitatory neurons from the awake condition. (A) Decoding error for visual stimuli as a function of SF from visual responses in the anesthetized (black) and awake conditions (red). The error represents the difference between the decoded and actual orientations. The dotted line indicates the chance level. ***p < 0.001 (anesthesia vs. awake). (B) Distributions of decoder probabilities for the visual responses in the anesthetized (left) and awake (right, control) conditions. The value of 0 on the SF and orientation axes indicates that the decoded stimulus matches the true stimulus. (C–D) Comparison of decoder probability profiles for the anesthetized and awake conditions. The profiles were extracted from (B) along the SF (C) and orientation axes (D), respectively. Under anesthesia, the probability of decoding at higher SFs was increased compared to the awake state, while the orientation did not change.

Next, to further investigate whether this degradation in information representation was a mere loss of precision or if it involved a specific representational bias, we analyzed the posterior probability generated by the decoder. The posterior probability represents the likelihood of each stimulus having been presented, given a specific pattern of observed neuronal activity. When we aligned the posterior probability centered on the true stimulus parameters (SF and direction) for each trial, this analysis revealed a systematic encoding bias under anesthesia (Fig. 3B). Specifically, when a high-SF stimulus was presented, the population response misrepresented it as if it were a lower-SF stimulus, evidenced by a significant shift in the peak of the posterior probability towards the lower frequencies (Fig. 3B–C). Notably, this bias was specific to SF, as the representation of stimulus direction remained unaffected by anesthesia (Fig. 3B and D). In contrast, decoding in the awake state consistently yielded high-fidelity representations for all stimuli (Fig. 3B–D). Taken together, these results strongly suggest that anesthesia selectively impairs the neural coding of high-SF visual information.

### Anesthetic Effects on Inhibitory Neuron Subtypes

We further investigated whether anesthesia-induced modulation varied across inhibitory neuron subtypes. We focused on SOM and PV neurons, the two major classes in mouse V1 that form monosynaptic inhibitory synapses onto excitatory neurons ^23–26^. SOM neurons were labeled using SOM-Cre;Ai14 mice (N = 5), and PV neurons with PV-Cre;Ai14 mice (N = 6).

Our analysis revealed that the SOM neurons displayed significantly lower preferred SFs under anesthesia compared to the awake condition (median: anesthetized 0.046 c/d, awake 0.068 c/d; p = 4×10^−5^, Kolmogorov–Smirnov test, n = 60 cells; Fig. 4A). The PV neurons also showed a significant decrease in preferred SFs under anesthesia (median: anesthetized 0.13 c/d, awake 0.22 c/d; p = 3×10^−7^, n = 33 cells; Fig. 4B). On average, preferred SFs decreased by 0.49 octaves after anesthesia in the SOM neurons (linear mixed-effects model, 95% CI [0.37 0.61], p = 2×10^−12^,), and by 0.69 octaves in the PV neurons (95% CI [0.53 0.85], p = 5×10^−12^). These results demonstrate that anesthesia generally modulates SF preference across these neuron subtypes.

**Figure 4.**
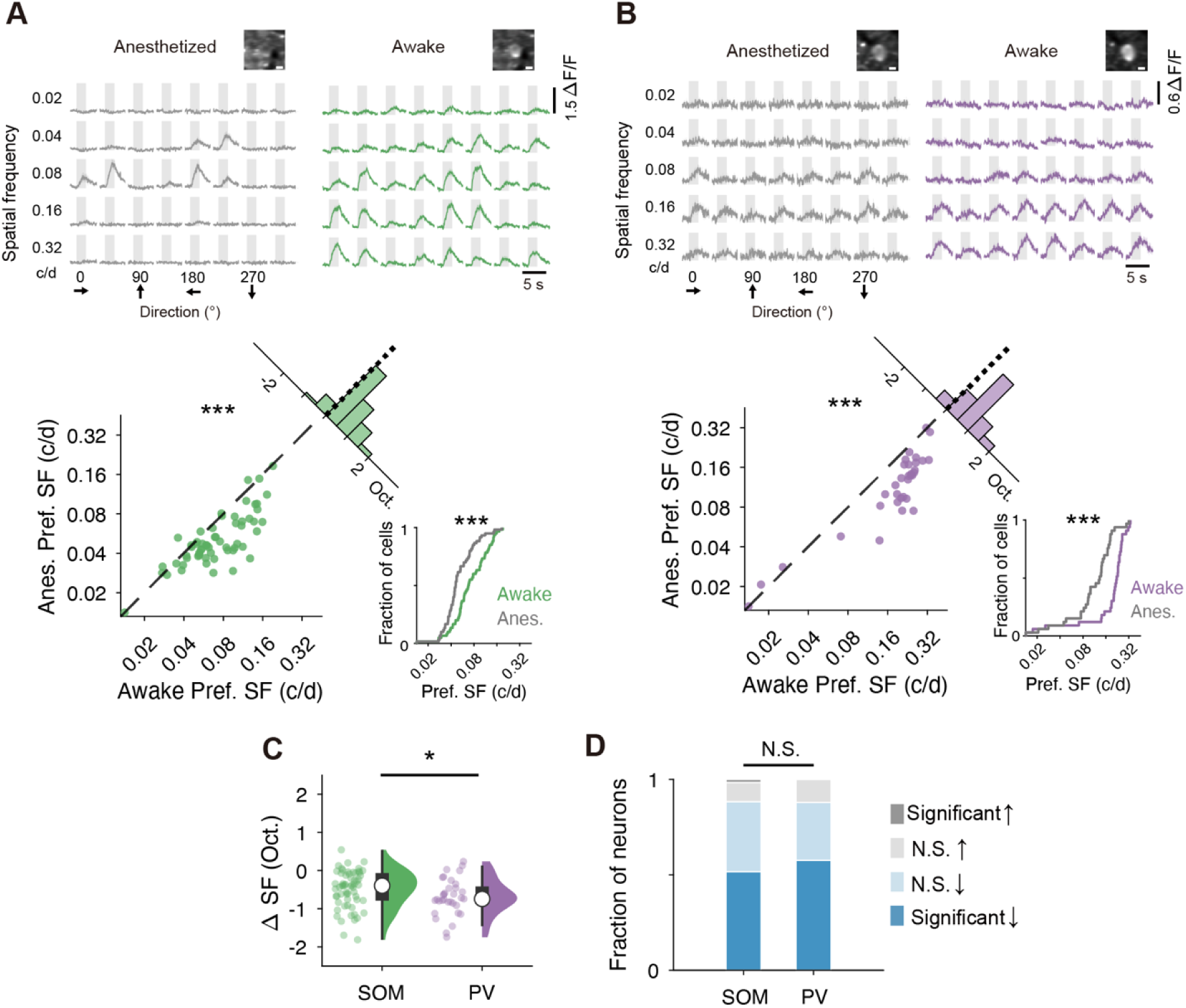
SOM and PV neurons show downward shifts of the preferred spatial frequency after anesthesia. (A) Comparison of preferred SFs between the anesthetized and awake states in the SOM neurons. Top, Representative visual responses. Bottom, Preferred SF of individual neurons in the anesthetized condition (y-axis) is plotted against that in the awake condition (x-axis) (n = 60 cells, N = 5 mice). Linear mixed model; ***p < 0.001. The corner histogram shows the distribution of the pairwise difference in preferred SF between the two conditions. Bottom right: cumulative distributions of preferred SFs in each condition. Kolmogorov–Smirnov test; ***p < 0.001. (B) Same plots for the PV neurons (n = 33 cells, N = 6 mice). (C) Box and raincloud plots comparing the anesthesia-induced changes in preferred SFs between the SOM and PV neurons. ΔSF indicates the difference in preferred SF (from the awake state to the anesthetized state) in each neuron. Wilcoxon signed-rank test; *p < 0.05. (D) Fractions of the SOM and PV neurons exhibiting a significantly decreased (blue), non-significantly (N.S.) changed (light blue or light gray), or significantly increased (gray) preferred SF under anesthesia. Comparison of fractions between the SOM and PV neurons. Chi-square test; N.S., not significant.

However, a direct comparison revealed that the SOM neurons showed a significantly smaller magnitude of anesthesia-induced shift in preferred SFs than the PV neurons (p = 0.046, Wilcoxon signed-rank test; Fig. 4C). The proportion of neurons exhibiting SF downshifts during anesthesia was not significantly different between the SOM and PV neurons (p = 0.29, Chi-square test, Fig. 4D). Additionally, the preferred SFs in the PV neurons were significantly higher than those in the SOM neurons, regardless of brain state (Fig. 4A–B, Fig. S3).

### Manner of Anesthesia-induced Preferred SF Shifts and Accompanying Changes in Visual Responsiveness

The anesthesia-induced downward shift in preferred SF could arise from two potential factors: (1) a reduced response to high SFs and/or an enhanced response to low SFs during anesthesia, or (2) a uniform shift of the entire SF tuning curve toward lower frequencies. To investigate these possibilities, we calculated the average population responses to all tested SFs in each neuron group. In excitatory neurons, the downward shift arose primarily from an increased response to low SF stimuli under anesthesia, rather than a decreased response to high SFs (Fig. 5A). Conversely, in inhibitory neurons, the downward shift was attributed to a specific reduction in responses to high SFs (Fig. 5B). Furthermore, we extended this analysis to the SOM and PV neurons. The downward shift was attributed to a specific reduction in responses to high SFs in the SOM neurons (Fig. 5C), while in the PV neurons, the entire SF tuning curve shifted toward lower frequencies (Fig. 5D). These findings suggest that anesthesia generally modulates SF preference across different neuron subtypes, but the underlying mechanisms driving this modulation are distinct.

**Figure 5.**
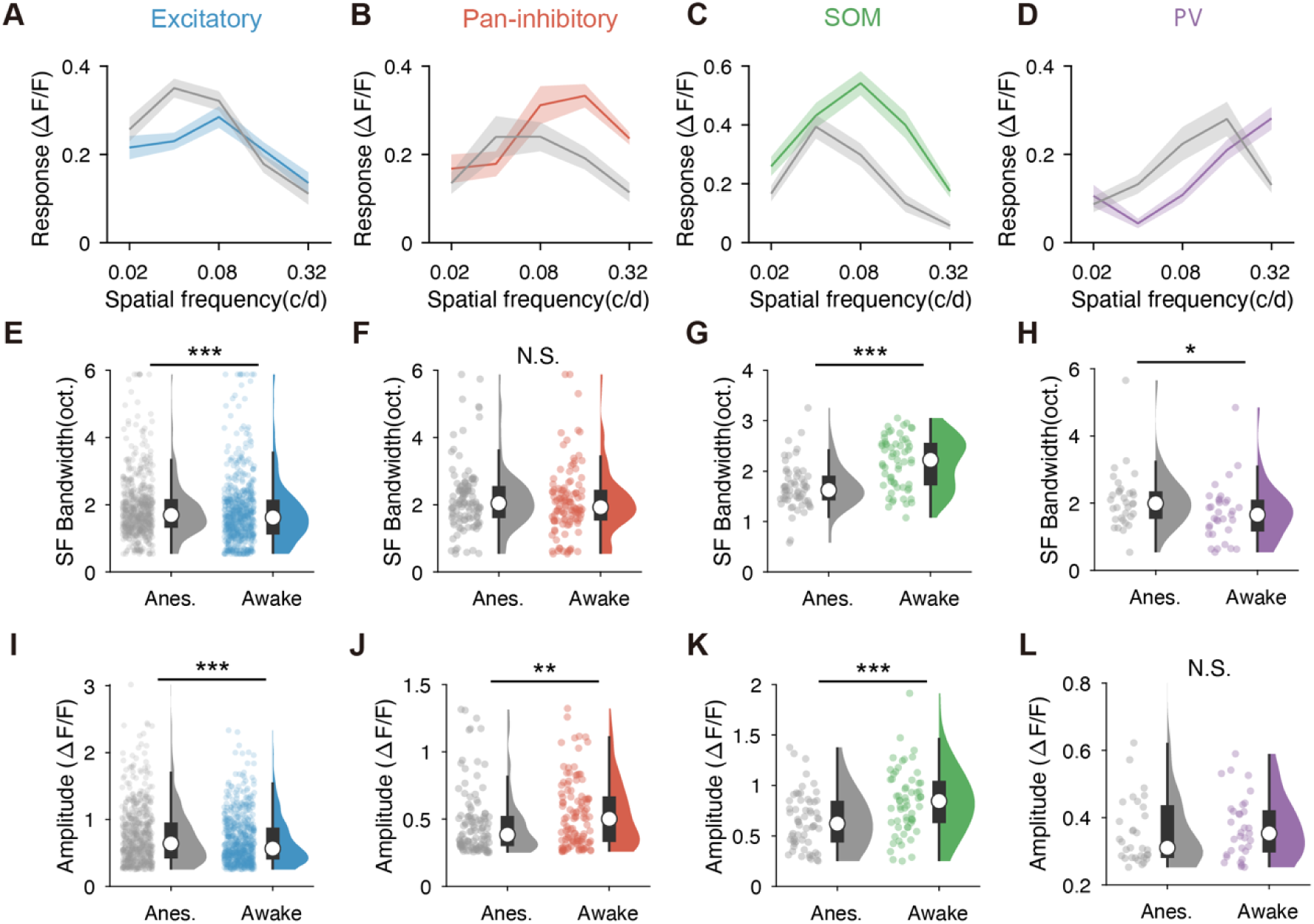
Manner of anesthesia-induced preferred SF shifts and accompanying changes in visual responsiveness. (A–D) Population-averaged SF tuning curves for excitatory neurons (A), pan-inhibitory neurons (B), the SOM neurons (C) and the PV neurons (D). Colored lines represent the awake condition, and gray lines represent the anesthetized condition. The error band represents the SEM. (E–H) Box and raincloud plots comparing the bandwidth of SF tuning curve for individual neurons in the anesthetized and awake conditions. (E) The excitatory neurons. (F) The pan-inhibitory neurons. (G) The SOM neurons. (H) The PV neurons. Wilcoxon signed-rank test; *p < 0.05, **p < 0.01, ***p < 0.001; N.S., not significant. (I–L) Similar to E–H, but for response amplitude.

The average population responses also implied anesthesia-induced changes in the sharpness of SF tuning and response amplitude between the anesthetized and awake states. To quantify these observations, we first compared the sharpness of SF selectivity between the two states. To this end, we calculated the full width at half maximum (FWHM) of the SF tuning curve in the optimal direction for individual neurons. The excitatory neurons showed a significantly broader SF tuning width in the anesthetized condition (median: 1.7 octaves) compared to wakefulness (1.6 octaves), indicating that SF selectivity became broader during anesthesia (p = 3×10^−6^, Wilcoxon signed-rank test; Fig. 5E). However, we found no significant differences in orientation or direction selectivity of the excitatory neurons between the anesthesia and awake states (p = 0.43 for orientation selectivity, p = 0.50 for direction selectivity; Fig. S4A and E), consistent with previous studies ^12,14,20^. These results suggest that anesthesia preferentially modifies SF tuning more than orientation and direction tuning in excitatory neurons.

The pan-inhibitory neurons displayed no significant differences in the sharpness of SF and direction selectivity between the awake and anesthetized conditions, but showed a significant difference in orientation selectivity (p = 0.46 for SF tuning, p = 1×10^−4^ for orientation selectivity, p = 0.54 for direction selectivity; Fig. 5F, Fig. S4B and F).

Interestingly, and in contrast to excitatory neurons, the SOM neurons exhibited significantly sharper SF tuning (i.e., a smaller FWHM) under anesthesia than during the awake state (median SF bandwidth: 1.6 octaves for anesthetized, 2.2 octaves for awake; p = 4×10^−6^; Fig. 5G). The SOM neurons also exhibited sharper selectivity for orientations and directions during anesthesia (p = 0.002 for orientation selectivity, p = 0.009 for direction selectivity; Fig. S4C and G). On the other hand, the PV neurons showed significantly broader SF tuning during anesthesia than during wakefulness (median SF bandwidth: 2.0 octaves for anesthetized, 1.7 octaves for awake; p = 0.02; Fig. 5H). The PV neurons showed no significant differences in the sharpness of orientation and direction selectivity between the two conditions (p = 0.99 for orientation selectivity; p = 0.37 for direction selectivity, Fig. S4D, H). These results demonstrate that the SOM neurons, but not the PV neurons, exhibited sharper visual tuning during anesthesia compared to wakefulness.

Regarding response amplitude, the responses to preferred visual stimuli during anesthesia showed a slight increase in the excitatory neurons compared to wakefulness (p = 1×10^−5^, Fig. 5I), whereas a decrease was observed in the pan-inhibitory neurons (p = 0.01, Fig. 5J). In the SOM neurons, visual response amplitude significantly decreased during anesthesia (p = 1.5×10^−5^, Fig. 5K), consistent with previous electrophysiological recordings ^15^. Conversely, the PV neurons showed no significant difference in the amplitude between the awake and the anesthetized conditions (p = 0.70, Fig. 5L), which also aligns with a previous analysis ^15^. These findings suggest that visual responses in SOM neurons are more sensitive to anesthesia than those in PV neurons. Overall, the changes in visual responses associated with the anesthesia-induced downshift in preferred SF appeared to be cell-type-specific.

### The Effect of Arousal Levels on Visual Responses

We finally investigated whether the downward shift of preferred SF is intrinsic to the anesthetic state, or if it also occurs during wakefulness in an arousal-dependent manner. To this end, we used pupil size as a proxy for arousal levels ^29,30^. Within awake sessions, trials were categorized as “low arousal” if the average pupil size during visual stimulation was in the lower 25% of all trials, and “high arousal” if in the upper 25% (Fig. 6A). Visual responses were analyzed from mice in which pupil size could be accurately analyzed (N = 6 mice, 216 cells).

**Figure 6.**
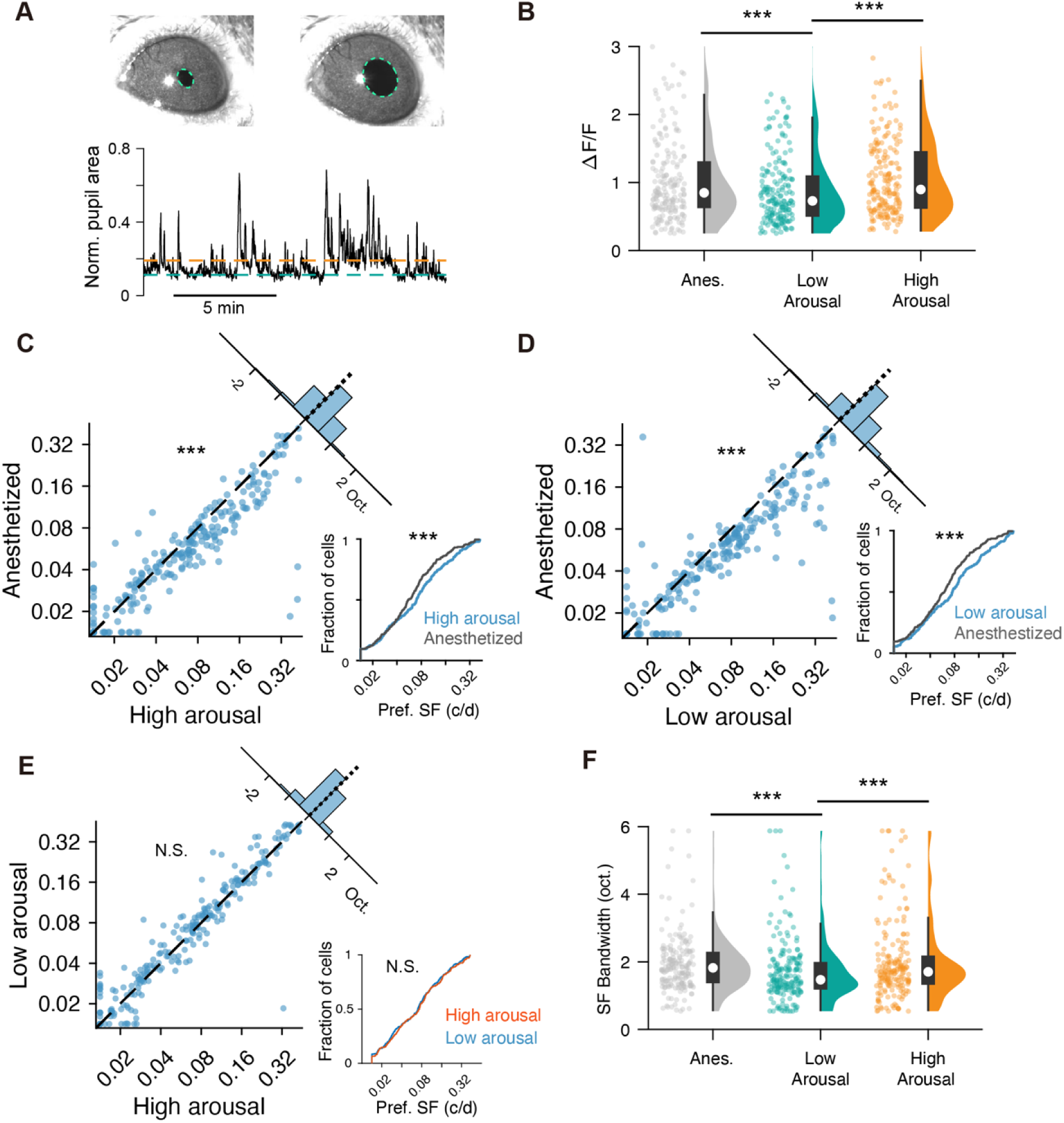
Comparison of visual responses between the anesthetic condition, and low-arousal and high-arousal states during wakefulness. (A) Estimation of arousal level from pupil area. Top: Example raw pupil trace. Bottom: Normalized pupil area trace. Dotted lines represent the 75th and 25th percentiles, used as thresholds for high- (orange) and low-arousal trials (green), respectively. (B) Box and raincloud plots comparing the visual response amplitude across the anesthetized, low-arousal, and high-arousal conditions (n = 216 cells, N = 6 mice). Wilcoxon signed-rank test with Bonferroni correction; ***p < 0.001. (C) Preferred SF of individual neurons in the anesthetized condition (y-axis) is plotted against that in the high arousal condition (x-axis). Linear mixed model; ***p < 0.001; N.S., not significant. The corner histogram shows the distribution of the pairwise difference in preferred SF between the two conditions. Bottom right: cumulative distributions of preferred SFs in each condition. Kolmogorov–Smirnov test; ***p < 0.001, N.S., not significant. (D–E) Similar to (C), but for comparison between the anesthetized and low-arousal conditions (D) and between the low-arousal and high-arousal conditions (E). (F) Similar to (B), but for SF tuning bandwidth.

The magnitude of visual responses in excitatory neurons was significantly smaller during low-arousal than during high-arousal trials, consistent with previous research (p = 3×10⁻⁷, Wilcoxon signed-rank test with Bonferroni correction; Fig. 6B) ^30^. The response magnitude under anesthesia was significantly larger than that during low-arousal trials (p = 1×10^-4^) but was comparable to that during high-arousal trials (p > 0.99). We then compared the preferred SFs in the high and low arousal states with those measured under the anesthetized condition. The preferred SFs were significantly higher in both low and high arousal states than during anesthesia (low arousal vs anesthesia: 0.32 octave, 95% CI [0.23,0.41], p = 2×10^-11^, linear mixed-effects model; high arousal vs anesthesia: 0.22 octave, 95% CI [0.14 0.30], p = 2×10^-17^, linear mixed-effects model, Fig. 6C–D). The preferred SFs did not differ significantly between the high and low arousal states, a finding consistent with previous research (p = 0.10, 95% CI [-0.01 0.10], linear mixed-effects model; p = 0.9, Kolmogorov–Smirnov test; Fig. 6E)^30^. Therefore, the downward shift in preferred SFs appears to be a specific effect of anesthesia. SF tuning sharpness was significantly lower (i.e., a larger FWHM) in the anesthetized state than in the low-arousal state (p = 2×10⁻⁵, Wilcoxon signed-rank test with Bonferroni correction, Fig. 6F). However, SF tuning sharpness was similar between the anesthetized and high-arousal states (p > 0.99), consistent with the corresponding changes in response magnitude.

## Discussion

By directly comparing visual responses in the same layer 2/3 neurons of V1 between the awake and anesthetized states, we demonstrate a systematic downward shift in preferred SF under isoflurane anesthesia. While this downward shift was observed in both excitatory and inhibitory neurons, the effect was more pronounced in inhibitory neurons, affecting both the SOM and PV interneuron subtypes. In line with these cellular-level observations, population decoding analyses revealed that the fidelity of representing high-SF stimuli was compromised under anesthesia, whereas the representation of low-SF stimuli remained robust.

In excitatory neurons, the anesthesia-induced downshift in preferred SF was accompanied by a broadening of SF tuning. Conversely, and consistent with previous reports, anesthesia did not alter the preference or tuning sharpness for orientation and direction in excitatory neurons ^12,14^.. Moreover, among the inhibitory neuron subtypes tested, anesthesia-induced changes in response gain and tuning sharpness were specific to SOM neurons but largely absent in PV neurons. Notably, optimal SFs were significantly higher in both low and high arousal awake states compared to the anesthetized condition. Taken together, these results indicate that anesthesia universally drives SF preference toward lower frequencies across neuronal subtypes, but that their magnitude and underlying mechanisms are highly subtype-specific.

### Anesthetic Modulation of SF Preference

Distinct anesthetic agents may induce various modulations in visual responses. Previous electrophysiological work in rat V1 has documented a decrease in visual responses to high SFs under isoflurane anesthesia compared to the awake state ^21^. Our data are consistent with this, demonstrating that isoflurane-induced downshifts in preferred SFs occur at the level of individual, identified neuron types in layer 2/3 of mouse V1. However, a previous study has shown no significant change in the preferred SFs of mouse V1 neurons between urethane-anesthetized and awake states ^14^. This suggests that the anesthesia-induced modulation of receptive field properties may depend on the choice of anesthetics. Another potential explanation for the different effects of anesthesia in the previous and current studies is that the earlier study did not restrict its analysis of preferred SFs to layer 2/3.

Ketamine is a dissociative anesthetic, and its primary molecular target is the NMDA receptor, which, despite some overlap, differs considerably from isoflurane’s primary target of GABA_A_ receptors ^5,22^. The effects on spontaneous cortical activity are also markedly different between ketamine and isoflurane ^31,32^. Ketamine induces a certain rhythm, originating in the retrosplenial cortex, which causes the dissociative state. Furthermore, under ketamine, spontaneous traveling waves propagate caudally across the cortex, entraining visual and parietal neurons similarly to stimulus-evoked waves observed during wakefulness ^31^. In contrast, this specific form of cortical entrainment is not observed under isoflurane anesthesia. Therefore, comparing awake and ketamine-anesthetized states may reveal different changes in SF preference than those observed between awake and isoflurane-anesthetized states.

### Potential Circuit Mechanisms for State-Dependent Changes in SF Preference

Clarifying the mechanisms that regulate SF preference across brain states remains challenging, as anesthesia’s effects are widespread and not limited to a specific neuronal population or specific circuits in V1. For instance, anesthetic-sensitive channels are expressed differentially across neuron subtypes^22^. Similarly, brain-state-dependent neuromodulators exert cell- or synapse-type-specific influences ^33^. These specific sensitivities to both anesthetics and neuromodulators likely represent a primary mechanism for modulating neural activity.

Consequently, these individual neuronal effects likely coalesce at the circuit level to produce the observed modulation of SF preference. The potential circuit factors can be broadly categorized into three areas: peripheral inputs, local neural circuits, and top-down inputs. The contribution of peripheral inputs seems unlikely, as visual responses in the lateral geniculate nucleus are similar between the anesthetized and awake states ^14^.

A second possibility is that local inhibition contributes to anesthesia-induced downward SF shifts. During wakefulness, excitatory neurons showed reduced responses to low SFs, shifting their preference toward high SFs. Although the SOM neuron responses were enhanced during wakefulness, this is unlikely to explain the suppression of low SF responses in excitatory neurons that receive direct inhibitory inputs from them. This is because the enhanced SOM neuron activity was specific to high SFs, while their responses to low SFs remained unchanged. Finally, top-down inputs from higher cortical areas are known to be considerably stronger during wakefulness compared to anesthesia^34,35^. Feedback neurons in the posteromedial area (PM), a higher visual area, preferentially respond to high SF stimuli and provide distinct excitatory inputs to V1 neurons that also prefer high SFs ^36–38^. During wakefulness, these feature-selective feedback inputs are likely significantly enhanced compared to anesthesia, potentially modulating the preferred SFs in V1.

In addition, another top-down pathway from the cingulate cortex preferentially activates vasoactive intestinal peptide-expressing (VIP) interneurons in V1 ^40^. Optogenetic activation of VIP neurons leads to an increase in the preferred SFs in excitatory neurons ^39^. Since VIP neurons directly inhibit SOM neurons, SOM neuron activity can serve as an inverse proxy for VIP activity. Our observation of elevated SOM neuron responses during wakefulness thus suggests low VIP neuron activity. Therefore, VIP neurons are unlikely to mediate the wakefulness-dependent increase in preferred SF.

To elucidate the mechanisms of anesthesia-induced changes in SF preference, further investigation is warranted. The evidence of anesthesia- or wakefulness-induced SF modulation could contribute to developing neuronal network models for SF representation in individual neurons ^41–43^, potentially leading to the elucidation of circuit mechanisms that cannot be revealed solely through physiological experiments.

### Differential Effects of Anesthesia on Visual Responses of SOM and PV Neurons

SOM and PV neurons are functionally distinct in their firing patterns, synaptic targets, and afferent input sources ^23–26^. SOM neurons primarily mediate feedback or lateral inhibition and are implicated in the formation of size tuning ^15,44^. In contrast, PV neurons predominantly drive feedforward inhibition and contribute to response gain regulation and the generation of high-frequency oscillations ^45–47^. Given that SOM and PV neurons constitute the major populations of inhibitory neurons, our results, indeed, suggest that their combined properties largely account for the findings observed in the pan-inhibitory population. We revealed that the preferred SF range of the SOM neurons was consistent with that of excitatory neurons and lower than that of the PV neurons, regardless of brain state (anesthetized or awake). Furthermore, we identified distinct patterns of anesthesia-induced SF shift between the SOM and PV neurons. Consistent with previous electrophysiological investigations ^15^, isoflurane anesthesia, relative to the awake state, diminished visual responses in the SOM neurons but not in the PV neurons. Additionally, we observed changes in the tuning sharpness for SF, orientation, and direction in the SOM neurons following anesthesia, whereas the PV neurons displayed almost no such alterations. These results demonstrate that the visual responsiveness of SOM neurons is profoundly modulated by the anesthetized state, which may help elucidate the distinct functional roles of these interneuron subtypes.

### Functional Significance

Our finding that isoflurane anesthesia impairs SF preference strongly suggests that the associated decline in sensory perception stems from a degradation of spatial resolution. This interpretation is consistent with prior work showing that V1 neuron size tuning, a key correlate of spatial resolution, is also impaired under anesthesia ^15,16^. Taken together, these findings suggest that anesthesia impairs spatial resolution at the level of single neurons in V1.

Furthermore, it remains an open question whether this reduction is specific to isoflurane or represents a more general feature of unconsciousness. To clarify this distinction, future experiments are warranted to examine the effects of other anesthetics and non-anesthetic states of unconsciousness like sleep ^1,5,31,32^. Ultimately, the consistent downward shift of preferred SFs under anesthesia not only reveals a potential mechanism for altered sensory perception but also highlights that high-SF tuning in the awake V1 is fundamental to perceiving fine visual detail.

## Methods

### Animals

All experimental procedures were performed in accordance with the guidelines of the Experimental Animal Committee of the National Institute for Physiological Sciences. Mice of either sex, aged 2–5 months (C57BL/6 background), were used. To visualize GABAergic, SOM-positive, and PV-positive interneurons, Vgat-IRES-Cre (B6J.129S6(FVB)-Slc32a1^tm2(cre)Lowl^/MwarJ, The Jackson Laboratory, stock #16962), SOM-IRES-Cre (B6N.Cg-Sst^tm2.1(cre)Zjh^/J; The Jackson Laboratory, stock #013044) and PV-IRES-Cre (B6;129P2-Pvalb^tm1(cre)Arbr^/J, The Jackson Laboratory, stock #008069) mice were respectively crossed with Ai14 tdTomato reporter mice (B6;129S6-Gt(ROSA)26Sor^tm14(CAG-tdTomato)Hze^/J, The Jackson Laboratory, stock #007908). The mice were housed under a standard visual environment (12/12 h light/dark cycles) with ad libitum access to food and water.

### Virus injection

For surgical procedures, mice were anesthetized with an intraperitoneal (ip) injection of a cocktail containing medetomidine (0.75 mg/kg; Domitor, Zenoaq), midazolam (4.0 mg/kg; Midazolam Sandoz, Sandoz), and butorphanol (5.0 mg/kg; Vetorphale, Meiji Seika). The animals were then head-fixed in a stereotaxic frame (SR-5M-HT, Narishige). A small craniotomy (< 0.4 mm diameter) was performed over the left visual cortex (2.7 mm lateral and 1.0 mm anterior to lambda). A beveled glass micropipette attached to a nanoinjector (Nanoject II, Drummond) was used to inject 400 nL of AAV1-Syn-jGCaMP7b-WPRE (Addgene) into layer 2/3 (400–500 μm from the dural surface). The viral solution was diluted with saline to a final titer of approximately 2–3×10^12^ genome copies/mL. Following the injection, the scalp was sutured. The mice were kept on a heating blanket until fully recovered from anesthesia.

### Craniotomy

A cranial window was implanted 10–12 days after the virus injection. Mice were anesthetized with an ip injection of a cocktail containing medetomidine (0.75 mg/kg), midazolam (4.0 mg/kg), and butorphanol (5.0 mg/kg). Throughout the surgery, body temperature was maintained with a heating blanket, and ofloxacin ophthalmic ointment was applied to the eyes to prevent corneal desiccation. To reduce inflammation and edema, mice were administered carprofen (6.0 mg/kg, ip), enrofloxacin (6.5 mg/kg, sc), and glycerol (0.15 ml/kg, ip).

The mice were mounted on a stereotaxic frame (SR-5M-HT, Narishige) with ear bars and a bite bar. The scalp overlying the parietal bones was resected, and the exposed skull was cleaned. A 3.5-mm-diameter craniotomy and durotomy were performed over the GCaMP7 expression site. A double-coverslip window was constructed by adhering a 3-mm-diameter coverslip (#3 thickness, Matsunami) to a 5-mm-diameter coverslip (#1 thickness, Matsunami) using a UV-curable optical adhesive (NOA61, Norland). This assembly was placed in the craniotomy. The edge of the window was then sealed to the surrounding skull first with cyanoacrylate adhesive (Aron Alpha A, Sankyo) and then reinforced with dental resin cement (Super-Bond C&B, Sun Medical). Finally, a custom-made head plate was attached to the skull using the same dental cement. The head plate was positioned to be as parallel as possible with the coverslip surface. Mice were monitored on a heating pad until they had fully recovered from anesthesia.

### Wide-Field Calcium Imaging

For wide-field Ca²⁺ imaging, visual stimuli were generated using PsychoPy ^49^ and presented on a 23-inch LCD monitor (FS2333, EIZO). The monitor was placed 20 cm from the mouse’s right eye, at a 45° angle to the right of the anteroposterior axis to be parallel to the right eye. Retinotopy mapping utilized a sweeping bar composed of a flickering black-and-white checkerboard pattern^48^. Each 15° square of the checkerboard inverted its contrast at 2 Hz. The bar was swept across the screen four times in each of the four cardinal directions (horizontal and vertical) at a speed of 10°/s. All stimuli were spherically corrected to account for the geometry of the planar monitor^36^.

Wide-field Ca^2+^ signals were recorded from head-fixed, awake mice. The cortical surface of the left hemisphere, including V1, was imaged at a 4-Hz frame rate using a Leica M165 FC stereomicroscope equipped with a cooled CCD camera (ORCA-ER, Hamamatsu Photonics) controlled by AQUACOSMOS software. Excitation light was filtered through a 450–490 nm bandpass filter, and emission signals were collected through a 500–550 nm bandpass filter. Analysis of the retinotopic mapping data was conducted using the ‘Visual Area Segmentation’ repository (https://snlc.github.io/ISI/; Marshel et al., 2011). The boundaries of V1 were identified from the resulting visual field sign map, and 500 × 500 µm regions of interest (ROIs) were defined within V1 for subsequent two-photon imaging.

### *In vivo* two-photon calcium imaging

GCaMP and tdTomato fluorescence signals were acquired using a two-photon microscope (A1MP, Nikon) equipped with a Ti:Sapphire laser (Mai Tai DeepSee, Spectra-Physics) and a 25× water-immersion objective (NA 1.10, MRD77220, Nikon). A custom-made light shield surrounded the objective to prevent light contamination from the stimulus monitor. Images (512 × 512 pixels; pixel size = 0.99 µm) were acquired at 15 frames/s using a resonant scanner. Both GCaMP7b and tdTomato were excited at 950 nm, and their respective emission signals were collected through bandpass filters (500/50 nm for GCaMP7b and 663/75 nm for tdTomato). Imaging was performed at depths ranging from 180–300 µm below the pial surface, corresponding to layer 2/3.

For all imaging sessions, mice were head-fixed under the microscope. Imaging was first performed in awake mice, followed by a session under light isoflurane anesthesia (0.6–0.8% in air). Each imaging session lasted approximately 50 min. During anesthetized recordings, body temperature was maintained at ∼37°C with a heating pad. In a subset of experiments, an additional awake session was conducted following a recovery period of at least 4 hours in the home cage. To compensate for slow axial drift between sessions, the imaging depth was re-adjusted using a custom ImageJ plugin by aligning the vasculature in the field of view to a reference stack recorded before the initial awake session. Data from sessions with axial motion artifacts exceeding 4 µm were excluded from further analysis.

### Visual stimulation

Visual stimuli were generated with PsychoPy ^49^and presented on a gamma-corrected 23-inch LCD monitor (FS2333, EIZO; 60 Hz refresh rate) positioned 20 cm from the mouse’s right eye. The display covered approximately 100° in azimuth and 70° in elevation of the visual field. The precise extent of this coverage was confirmed at the start of each experiment by mapping neuronal responses to a flickering checkerboard stimulus.

The stimuli consisted of full-screen, sine-wave drifting gratings (100% contrast; 100 cd/m² mean luminance) presented to the right eye. These gratings utilized six spatial frequencies (SFs) (0.02–0.64 cycles/degree, in one-octave steps) and eight directions of motion (0–315°, in 45° steps), at a temporal frequency of 2 Hz. The stimulus set also included a blank condition (a gray screen at mean luminance) and a 2 Hz full-field flicker. A total of 50 unique stimuli were presented in a pseudo-random order, with 10 repetitions per imaging session. Each trial consisted of a 2-s stimulus presentation, followed by a 4-s inter-trial interval showing a gray screen.

### Estimation of visual tuning preferences

Raw two-photon imaging data were motion-corrected, and regions of interest (ROIs) corresponding to individual neurons were identified using Suite2p^51^. For mice expressing both GCaMP and tdTomato, the structurally-expressed tdTomato fluorescence (red channel) was used for motion correction, while ROIs were detected from the functionally-driven GCaMP signal in the green channel. Putative non-neuronal ROIs were manually curated and excluded in the Suite2p GUI.

To correct for neuropil contamination, the neuropil signal for each ROI was defined as the average fluorescence within a circular annulus (inner edge 3 µm from the ROI boundary; width 5 µm). The fluorescence time series of neurons (*F_soma_*) was then corrected by subtracting the scaled neuropil signal (*F_neuropil_*) according to the formula:

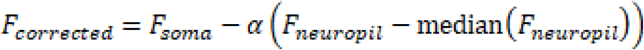

for each neuron was estimated using robust linear regression between its cell fluorescence and the corresponding neuropil signal. If ***α*** exceeded 1, ***α*** was set to 0.93 (median value across all cells). The relative change in fluorescence (ΔF/F) was calculated for each trial as (Fcorrected −F0)/F0, where F0 is the mean fluorescence during the 1.3-s period immediately preceding stimulus onset.

The response amplitude for each stimulus condition was computed as the average ΔF/F across 10 repetitions. A neuron was classified as visually responsive if it met two criteria: (1) its activity during periods of visual stimulation was significantly different from the blank condition (one-way ANOVA, p<0.01), and (2) its trial-averaged response to the preferred stimulus exceeded a threshold of 0.25 ΔF/F. The number of responsive cells in each neuron subtype is summarized in Table S1. For the analysis, we selected neurons responsive to visual stimuli in both the anesthetized and awake states.

The direction and SF tuning properties for individual neurons were characterized by fitting their trial-averaged responses to the product of a double Gaussian in direction space and a Gaussian in SF space. The response ***R*** is described as:

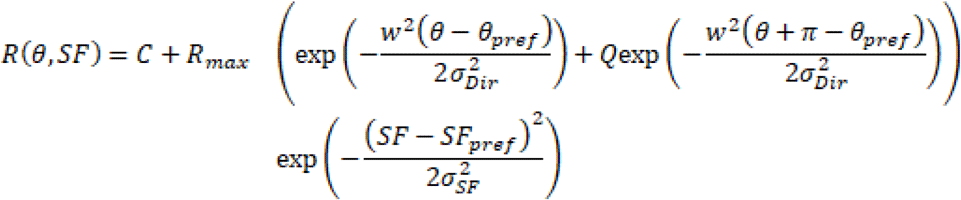

***w*(*θ*)** wrap angles onto the interval between. ***0*** and ***π***:

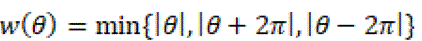

where **R(*θ, SF*)** is the trial-averaged ΔF/F for the stimulus with direction ***θ***and spatial frequency ***SF***. This model assumed that the peaks of the two Gaussians were 180° apart. ***θ_prof_*** is the preferred direction, **<u>σ*_Dir_*</u>** is the width of the direction tuning curve, ***Q*** is the relative magnitude of the response in the null direction, ***SF_pref_*** is the preferred SF, **σ*_SF_*** is the width of the SF tuning curve, ***R_max_*** is the response to the preferred SF and direction, and ***C*** is a constant amplitude offset. The values of the fit parameters were determined by minimizing the squared error using the lsqcurvefit function in MATLAB (R2020b, MathWorks). Fits with a low coefficient of determination (R^2^<0.3) were excluded from the SF analysis.

To assess the reliability of the estimated tuning parameters, a bootstrapping procedure was performed. For each neuron, we generated 1000 surrogate datasets by randomly resampling (with replacement) the 10 trials for each grating stimulus. The tuning curve was re-fit to each surrogate dataset, yielding a distribution of tuning parameters, from which 95% confidence intervals (CIs) were derived. A neuron was considered to exhibit a significant difference in a tuning parameter (e.g., preferred SF) between the anesthetized and awake conditions if the respective 95% CIs did not overlap.

The sharpness of the SF tuning was assessed by the full width at half maximum (FWHM) of the tuning curve at the optimal direction. The orientation selectivity index (OSI), computed as follows:

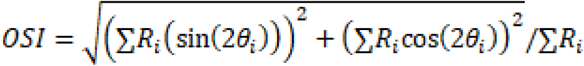

Where ***R_i_*** is the response to the ***i^th^*** direction with the optimal SF.

The direction selectivity index (DSI) was calculated as follows:

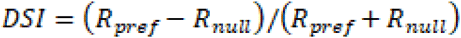

where ***R_prof_*** is the response in the preferred direction, and ***R_null_*** is the response in the opposite direction.

### Bayesian decoding

To examine how anesthesia influenced the representation of visual information in the neural population activity, we implemented a Bayesian decoding analysis based on the framework of Jazayeri and Movshon ^50^. This method estimates the probability that a given population response, *r* = {*r_1_, r_2_, …, r_N_* }, was generated by a stimulus, *s*, from the set of all possible stimuli, *S*.

We assumed that the neural response, *r_i_*, from each neuron, *i*, on a given trial was drawn from an independent Poisson distribution with a mean response defined by its tuning curve, ***f_i_*(*s*)**. For each neuron ***i***, we calculated the tuning function ***f_i_*(*x*)** as the mean response across trials to stimulus ***s***. Assuming a uniform prior probability for all stimuli, the log-likelihood of a stimulus ***s*** given a population response ***r*** can be computed as a weighted sum of the responses from all ***N*** neurons:

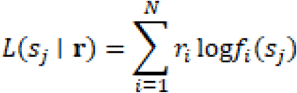

The posterior probability of the stimulus, ***P*(*s_j_*|*r*)**, was calculated by normalizing the likelihood across all possible stimuli:

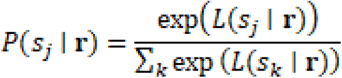

For each trial, the posterior probability distribution over the stimulus set was computed. The decoded stimulus was defined as the one with the highest posterior probability (the maximum a posteriori, or MAP, estimate). Decoding error was quantified as the absolute difference between the decoded stimulus orientation and the true stimulus orientation. We used level-one-out cross-validation to decode the awake state response with the awake state tuning curve. Trials that had been decoded were excluded from calculating the tuning curve.

To further assess how anesthesia influenced the representation of visual information, we also examined the full posterior probability distribution. Unlike the single MAP estimate, the full distribution provides more information about the representational landscape, revealing changes such as bias or systematic shifts. Before MAP estimation, these probabilities were normalized to sum to 1 over all possible stimuli. After computing per-trial posteriors, they were aligned to the true stimulus by centering them on that trial’s stimulus value. Then, the aligned posterior probabilities were averaged across trials within each state, and the resulting distributions were compared between the awake and anesthetized conditions.

### Arousal modulation

The animal’s arousal state was monitored by tracking pupil diameter, following established methods ^29,30^. To avoid interfering with visual stimulation of the right eye, the contralateral (left) eye was illuminated with an infrared (IR) LED, and videos of the eye were recorded at 15 Hz using a monochrome camera (MQ003MG-CM, XIMEA) equipped with a macro zoom lens (MLM-3XMP, COMPUTAR) and a long-pass filter (>830 nm; LP830, AZURE). Video acquisition was synchronized with frame acquisition triggers from the two-photon microscope using custom Python software.

Pupil size and position were extracted from the recorded videos offline using Facemap (https://github.com/MouseLand/facemap). For each session, a region of interest (ROI) encompassing the pupil was manually drawn. To account for ambient light fluctuations, the minimum pixel value within the ROI was subtracted from all pixels on a frame-by-frame basis. A brightness threshold was then manually set to isolate dark pupil pixels. The pupil’s area and center were calculated by fitting a 2D Gaussian to the covariance of these threshold pixels.

The resulting pupil area trace was filtered to remove blink-related artifacts. Specifically, a median filter (2-s window) was applied, and any raw data point that deviated from the median by more than 0.5 standard deviations was replaced with the median-filtered value. The cleaned trace was then normalized to the maximum pupil area observed within that session.

For subsequent analysis, trials were categorized into low and high arousal states based on the session-wide pupil area distribution. ‘Low arousal’ corresponded to pupil areas below the 25th percentile, while ‘high arousal’ corresponded to areas above the 75th percentile. The arousal level for a given trial was defined as the average normalized pupil area during that trial’s stimulus presentation. Trials were subsequently categorized as ‘low arousal’ or ‘high arousal’ if this average value fell into the corresponding percentile range.

Finally, to investigate the effect of arousal on sensory processing, SF and direction tuning curves were fitted separately to neuronal responses during low and high arousal trials. For SF tuning analysis, a stricter goodness-of-fit threshold (R^2^>0.4) was used, to account for the relatively low response amplitudes observed in low arousal trials.

### Statistical analysis

All quantitative and statistical analyses were performed using custom scripts written in MATLAB (R2020b, MathWorks). Data are presented as mean ± standard deviation (SD). In box plots, the central mark indicates the median, the bottom and top edges of the box indicate the 25th and 75th percentiles, and the whiskers extend to the most extreme data points not considered outliers. Outliers were defined as data points falling more than 1.5 times the interquartile range (IQR) beyond the box edges.

Statistical tests were chosen based on the data structure. The Wilcoxon signed-rank test was used for paired comparisons, while the Wilcoxon rank-sum test or the two-sample Kolmogorov-Smirnov test was used for unpaired group comparisons. To compare the distributions of preferred orientations, circular two-sample Kuiper tests were performed using the CircStat toolbox for Matlab^52^. For multiple comparisons, a Bonferroni correction was applied to adjust the significance threshold. Correlations were assessed using Pearson’s correlation coefficient. The chi-squared (χ^2^) test was used for comparing proportions between two groups. Unless otherwise stated, all statistical tests were two-sided, and a p-value of < 0.05 was considered statistically significant.

To account for the non-independence of data from the same animal, linear relationships between preferred SFs in different brain states were tested using linear mixed-effects models (Statistics and Machine Learning Toolbox, MATLAB). In these models, individual animal identity was included as a random effect.

## Data avxailability

All data and code generated for producing the figures in this manuscript will be deposited and available at https://github.com/jiahao-wu34/Brain_states_spatial_frequency upon publication of the Version of Record.

## Acknowledgements

We thank Dr. G. Okazawa for critically reading this manuscript and members of the Yoshimura laboratory for helpful discussions. We also thank M. Takagi, K. Ishigami, S. Omura, and M. Higa for technical assistance. YY was supported by JSPS Grant-in-Aid for Scientific Research (B) (21H026000, 24K02140), and Grant-in-Aid for Challenging Research (Pioneering) (23K17409). NT was supported by National Health Medical Research Council (GNT1183280, GNT2037172), Australian Research Council (DP240102680), JSPS of Science Grant-in-Aid for Transformative Research Areas (A) (23H04829, 23H04830) and JST Moonshot R&D Grant (JPMJMS2295-14), Theoretical Sciences Visiting Program, Okinawa Institute of Science and Technology.

## Author contributions

J.W., T.Y. and Y.Y. designed research; J.W. performed research; J.W. and T.Y. analyzed data with input from K.R. and N.T.; and J.W. and Y.Y. wrote the original draft, and all authors contributed to the final version.

