## Supplementary Figures for "Anesthesia Lowers Spatial Frequency Preference in the Primary Visual Cortex"

**Supplementary Information**

**Figure S1–S4**

**Table S1**

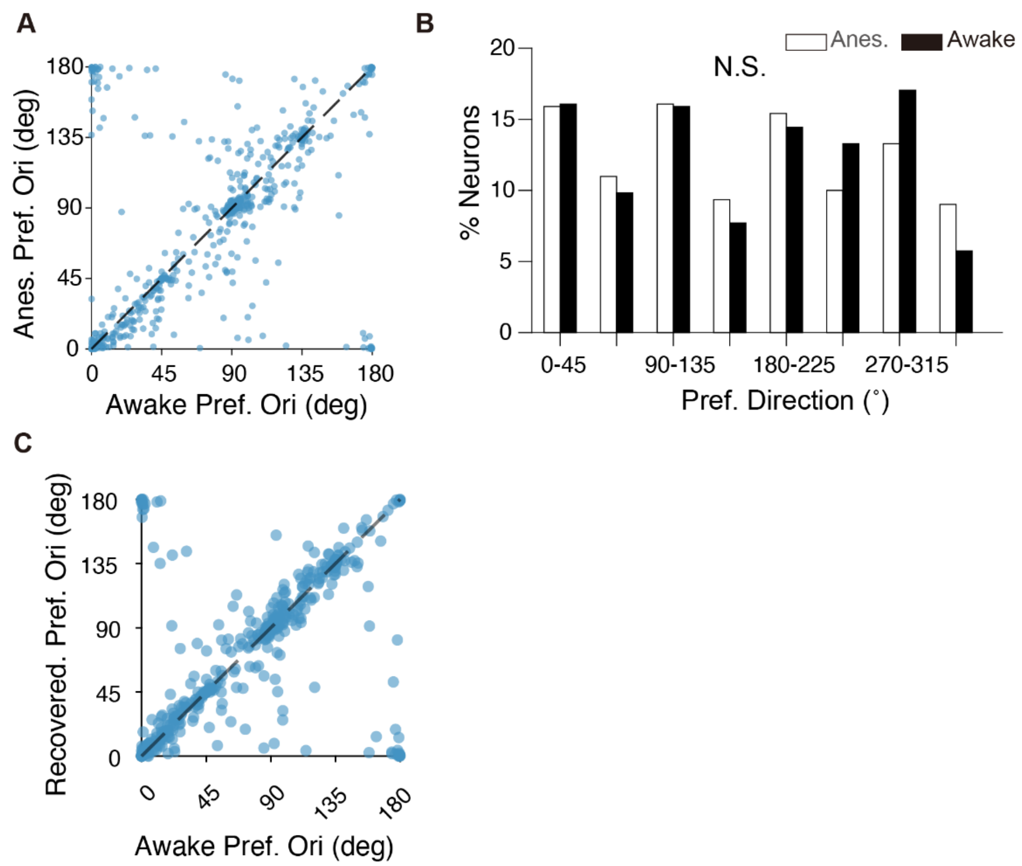


**Supplement Figure 1. Comparison of preferred orientation in excitatory neurons between the anesthetic and awake conditions.**

(A) Preferred orientation of individual neurons in the anesthetized condition (y-axis) is plotted against that in the awake condition (x-axis, n = 610 cells, N = 9 mice).

(B) Distribution of preferred orientations in the anesthetized and awake conditions. The Kuiper two-sample test; N.S., not significant.

(C) Preferred orientation of individual neurons in the recovery condition from the anesthesia ($2^{nd}$ awake, y-axis) is plotted against that in the $1^{st}$ awake condition (x-axis) (n = 376 cells, N = 5 mice).

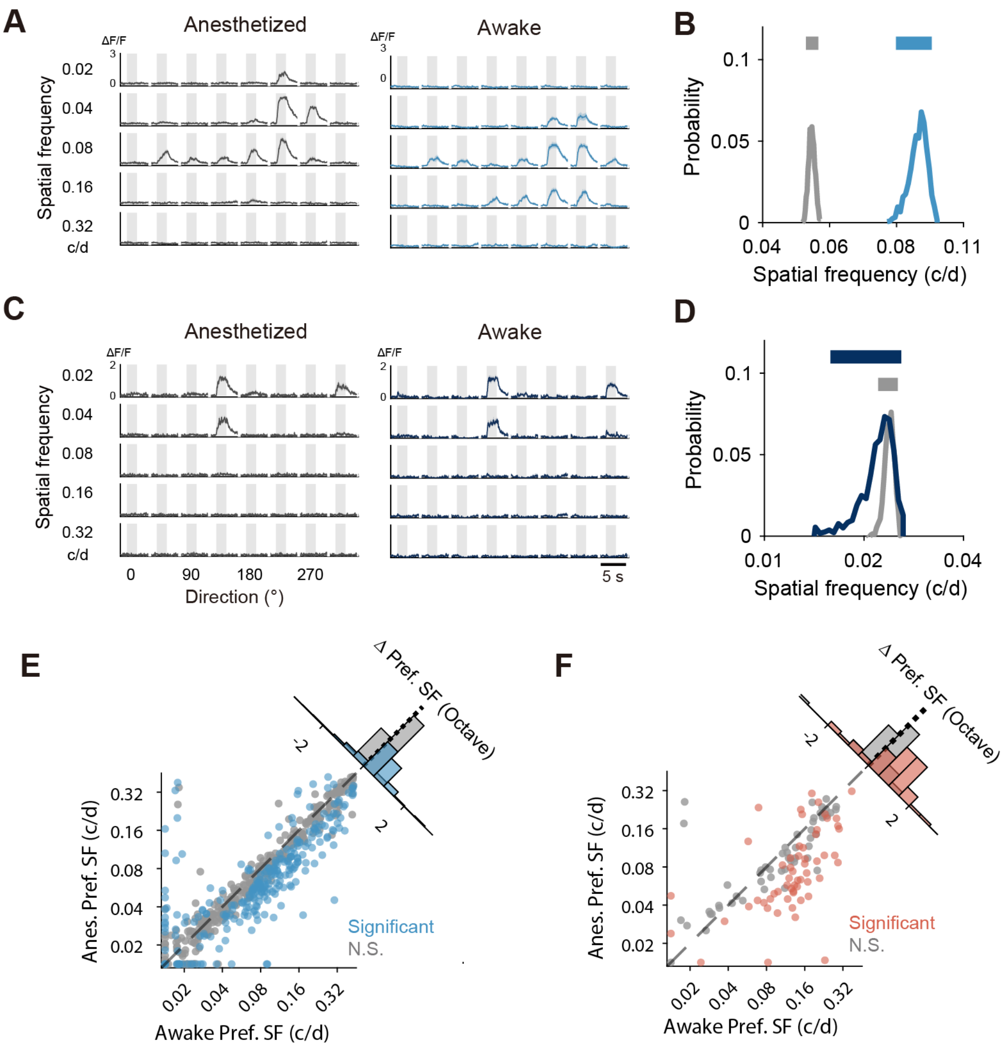


**Supplement Figure 2. Estimation of significantly SF shifted neurons under anesthesia by bootstrapping analysis.**

(A) Example visual responses of an excitatory neuron in the anesthetized (left) and the awake (right) conditions (mean ± SEM). Responses (ΔF/F) were evoked by visual stimuli with various spatial frequencies (1-octave steps, shown on the left) and directions (45°-steps, shown at the bottom). Each gray box indicates the stimulus presentation period (2 s).

(B) Distribution of preferred SFs estimated by a bootstrapping method in anesthetized (gray) and awake (blue) conditions for the neuron shown in (A). Bars above the histograms represent the 95% confidence intervals. This neuron was classified as an SF-shifted neuron.

(C–D) Similar to (A) and (B), but for a neuron that showed no significant shift in preferred SF.

(E–F) Comparison of preferred SFs between the anesthetized and awake conditions for excitatory neurons (E) and inhibitory neurons (F). The same plot as Fig. 2A and C, but neurons were categorized as shifted (color) or unshifted (gray) based on the bootstrap analysis. The corner histogram shows the distribution of the difference in preferred SFs (octave) between the two conditions.

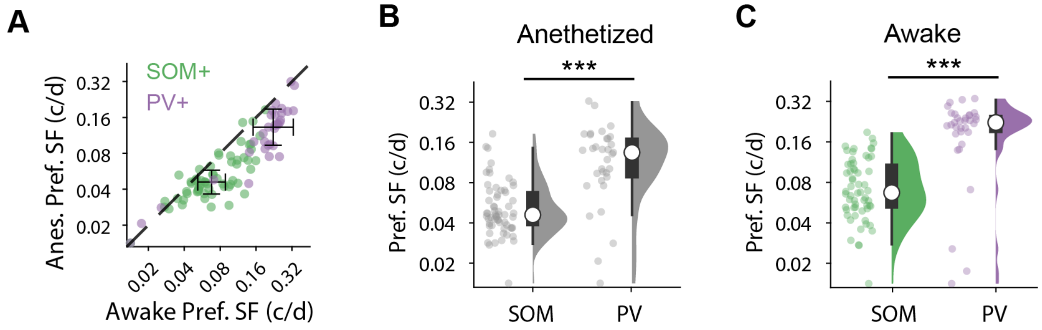


**Supplement Figure 3. The preferred SFs are higher in the PV neurons than SOM neurons.**

(A) Comparison of preferred SFs between the anesthetized and awake conditions for the SOM (green) and PV (purple) neurons. The plot superimposes data from Fig. 4A and B.

(B–C) Box and raincloud plots comparing the preferred SFs in the anesthetized (B) or awake conditions (C) between the SOM and PV neurons. Wilcoxon rank-sum test;. ***P < 0.001.

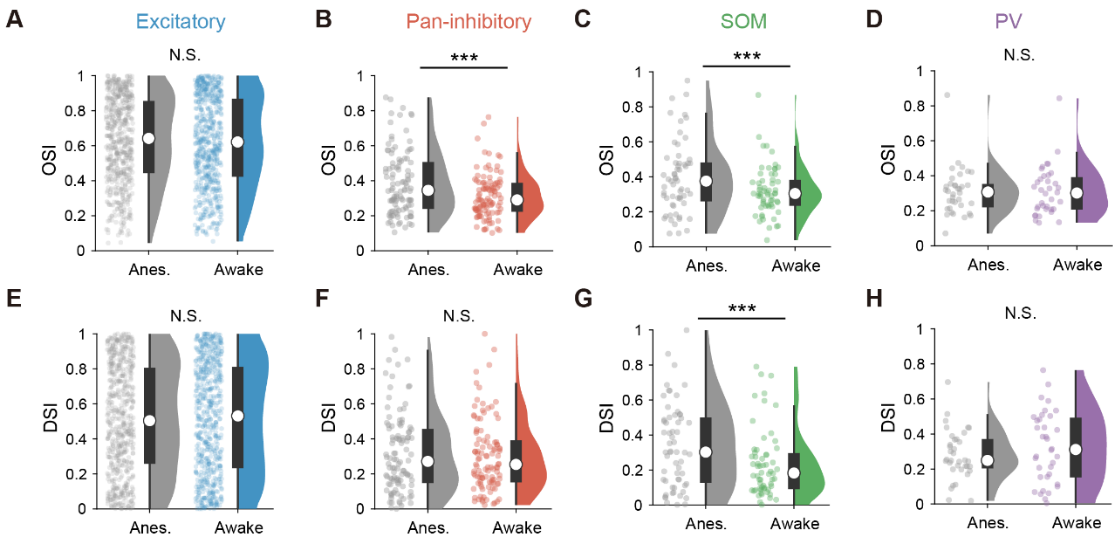


**Supplement Figure 4. Comparison of orientation and direction selectivity between anesthetized and awake conditions.**

(A–D) Box and raincloud plots comparing the orientation selectivity index (OSI) calculated from the responses to preferred SFs in individual cells in the anesthetized (gray) and awake (colored) conditions. (A) Excitatory neurons (n = 610 cells, N = 9 mice). (B) Pan-inhibitory neurons (n = 111 cells, N = 9 mice). (C) the SOM neurons (n = 60 cells, N = 5 mice). (D) the PV neurons (n = 33 cells, N = 6 mice).

(E–H) Same plot as (A–D), but for the direction selectivity index (DSI). Wilcoxon signed-rank test; ***$p < 0.001$. N. S. is not significant.

**Table S1. The number of visually responsive cells.**

| Cell type | N (animals) | States | Responsive/Total | Proportion |
|---|---|---|---|---|
| **Excitatory neurons** | 9 | Anesthesia | 916/2029 | 45.1% |
| | | Awake | 803/2029 | 39.6% |
| | | Both | 610/2029 | 30.5% |
| **Pan-inhibitory neurons** | 9 | Anesthesia | 174/446 | 39.0% |
| | | Awake | 195/446 | 43.7% |
| | | Both | 111/446 | 27.4% |
| **SOM neurons** | 5 | Anesthesia | 64/89 | 71.9% |
| | | Awake | 74/89 | 83.1% |
| | | Both | 60/89 | 68.5% |
| **PV neurons** | 6 | Anesthesia | 53/116 | 45.7% |
| | | Awake | 55/116 | 47.4% |
| | | Both | 33/116 | 36% |
